# Cellular aging in vitro recapitulates multi-tissue epigenetic aging in vivo

**DOI:** 10.1101/2020.09.02.280073

**Authors:** Christopher Minteer, Marco Morselli, Margarita Meer, Jian Cao, Albert Higgins-Chen, Sabine Lang, Matteo Pellegrini, Qin Yan, Morgan Levine

**Affiliations:** Department of Pathology, Yale School of Medicine, New Haven, CT, USA 06519; Department of Molecular, Cell, and Developmental Biology, University of California Los Angeles, Los Angeles, CA, USA 90095; Rutgers Cancer Institute of New Jersey, New Brunswick, NJ, USA 08901

**Keywords:** Aging, DNA methylation, epigenome, differentiation, caloric restriction, iPSC, replicative senescent

## Abstract

Aging is known to elicit dramatic changes to DNA methylation (DNAm), although the causes and consequences of such alterations are not clear and require further exploration. Our ability to experimentally uncover mechanisms of epigenetic aging will be greatly enhanced by our ability to study and manipulate these changes using *in vitro* models. However, it remains unclear whether the changes elicited by cells in culture can serve as a model of what is observed in aging tissues *in vivo*. To test this, we serially passaged mouse embryonic fibroblasts (MEFs) and assessed changes in DNAm at each time-point via RRBS. By developing a measure that tracked cellular aging *in vitro*, we tested whether it tracked physiological aging in various mouse tissues and whether anti-aging interventions modulate this measure. Our measure, termed DNAmCULTURE, was shown to strongly increase with age when examined in multiple tissues (liver, lung, kidney, blood, and adipose). As a control, we confirmed that the measure was not a marker of cellular senescence, suggesting that it reflects a distinct yet progressive cellular aging phenomena that can be induced *in vitro*. Furthermore, we demonstrated slower epigenetic aging in animals undergoing caloric restriction and a resetting of our measure in lung and kidney fibroblasts when re-programmed to iPSCs. Enrichment and clustering analysis implicated SUZ12, EED and Polycomb group (PcG) factors as potentially important chromatin regulators in translational culture aging phenotypes. Overall, this study supports the concept that physiologically relevant aging changes can be induced *in vitro* and moving forward, used to uncover mechanistic insights into epigenetic aging.

## iii. Introduction

Aging is characterized by a progressive decline in cell, tissue and organ integrity that manifests as age-related diseases and ultimately death [1,2]. Telomere attrition [3], cellular senescence [4-6], DNA damage [7], stem cell exhaustion [8] and epigenetic modifications [9] represent just a few molecular and cellular features of the aging process. While these hallmarks have been extensively investigated, their interactions, causes, and the resulting emergence that leads to the failure of the organism is not well characterized. Epigenetic alterations in aging—specifically alterations in DNA methylation (DNAm)—is a clear example of a hallmark which has been widely studied, but lacks a conceptual mechanistic framework linking its causes and consequences to other hallmarks or physiological manifestations with aging.

DNA methylation (DNAm) refers to the addition of a methyl group (CH3) to a CpG dinucleotide (5;—C—phosphate—G—3’). In most cases, DNAm is associated with transcriptional repression via its effect on chromatin accessibility, and is thought to control a number of cellular properties, including differentiation [10], replication [11], X-inactivation [12], stress response [13], and genomic imprinting [14]. Initially, *de novo* methyltransferases establish methylation patterns that are necessary for organismal development [15,16]. These patterns are then modulated by maintenance methyltransferases over the course of the lifespan [17,18] during which, subtle changes can dramatically alter promoter function [19-21] and distal regulatory elements [22,23]. Changes in DNAm with aging were first reported more than three decades ago and now occupy a major field in aging research [24]. These changes paint a picture characterized by a gain of DNAm at gene promotors and loss of global DNAm, representing trends towards hypomethylation in intergenic regions associated with dispersed retrotransposons, heterochromatic DNA repeats, and endogenous retroviral elements [25]. Given the predictability of these age-related changes, researchers began applying machine learning techniques to develop age predictors from DNAm that could serve as biomarkers of aging. To date, these so called “epigenetic clocks” have been applied in a plethora of tissues across diverse mammalian species and have been shown to be predictive of lifespan and health span, above and beyond chorological age [26-30]. Although exciting, the mechanistic underpinnings and drivers of epigenetic clocks are relatively unknown, limiting the conclusions that can be drawn.

Our lack of mechanistic understanding when it comes to epigenetic clocks likely stems from the fact that these models have been almost exclusively applied to *in vivo* and *ex vivo* blood and tissue samples in humans (and more recently in other mammals) for which experimental investigation is limited. Thus, we hypothesize that the use of culture models coupled with physiologically relevant tissue samples may greatly facilitate mechanistic discovery.

Culture aging within the context of cellular biology is extensively examined, presenting a model to study mechanisms of epigenetic aging [31,32]. From the time Hayflick proposed the theory now known as the Hayflick limit [33], many studies have contributed to characterizing exhaustive passaging, providing robust and well characterized culture models that can be used to determine the extent culture aging recapitulates physiological aging [34-38]. However, none have applied systems-level measures to directly demonstrate whether changes that can be induced in culture mimic what happens with aging in the organism. Thus, the aims of this paper were i) to better characterize the culture aging phenomena by generating a clock based on DNA methylation changes *in vitro*, ii) test whether such culture models of aging capture a physiologically relevant signal, and iii) use this data as a first step towards elucidating mechanisms of aging. Overall, the results from this study sets the foundation for using culture aging epigenetic models as a translational bridge to *in vivo* biomarker studies.

## Results

### Developing a measure of culture aging using DNAm

To explore culture aging, understand its association with the methylome and determine the extent culture phenotypes recapitulate physiological aging, we derived a primary mouse embryonic fibroblast culture system that was exhaustively passaged to produce longitudinal DNAm samples (Figure 1A, Supplemental Figures 1A-D). We selected mouse embryonic fibroblasts (MEFs) as our model, given their accelerated aging phenotype after relatively few passages (5-7) under normoxic (20%) conditions [35,37]. This accelerated aging is hypothesized to occur from extrinsic factors, like oxygen toxicity, rather than intrinsic factors like telomere shortening [31]. It is also a distinct phenotype in contrast to MEFs grown under physiological conditions of 3% oxygen, which do not senesce. Given that genotoxic stress is known to modulate the methylome [39-41], we reasoned that this model will enable us to capture the known murine sensitivity to oxidative damage using DNAm from serially passaged MEFs under normoxia.

**Figure 1:**
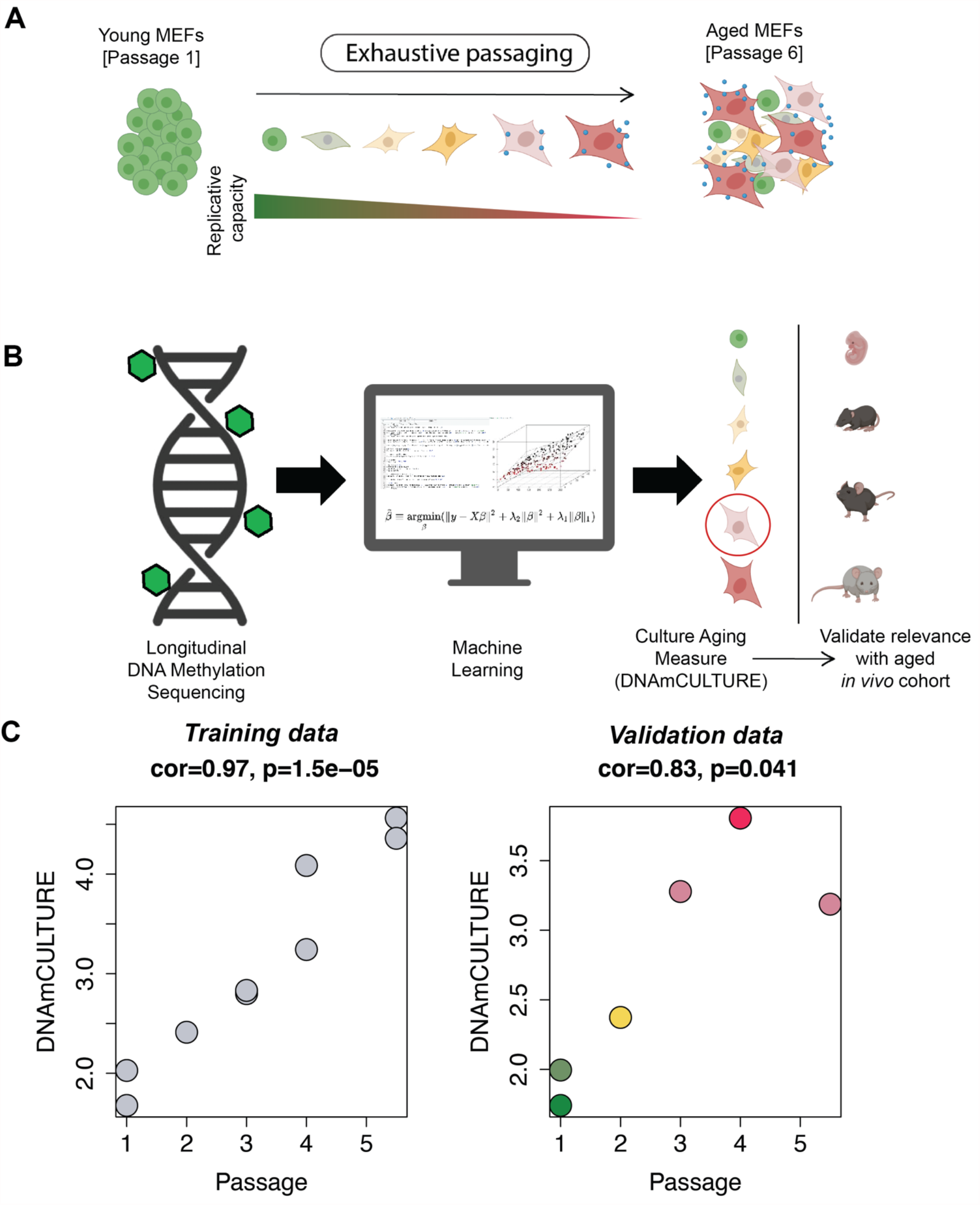
Development of a DNAm culture aging measure (DNAmCULTURE) in mouse embryonic fibroblasts. (A) Schematic displaying exhaustive culturing of mouse embryonic fibroblasts under normoxia (20% O2) produces terminally arrested cellular states with progressively reduced replicative capacity. (B) Workflow demonstrating supervised machine learning computation approach (elastic net penalized regression) successfully produced a measure of culture aging from longitudinal reduced represented bisulfide sequencing (RRBS) DNA methylation data, where it was then was tested for physiological relevance in an aged *in vivo* cohort. (C) Training (MEF1 and MEF2) and validation (MEF3) cell lines used to develop DNAmCULTURE, with shading representing predicted culture age in validation data. Note, passage 5 and 6 were combined due to low DNA content prior to RRBS sequencing and are represented throughout as passage 5.5. Passage correlations and statistical significance was determined using Pearson correlations.

DNAm was assessed at each passage in three biological replicates via reduced representation bisulfite sequencing (RRBS) with the goal of utilizing machine learning techniques to reduce the highly dimensional DNAm data into a single meaningful measure that increases as a function of time in culture (Figure 1B). The primary data used to train the culture measure, termed DNAmCULTURE, was obtained from passages 1-6 of the culture MEF system. Of the three MEF cell lines, two were used in training and the third was used for validation. In both cases, passages 5 and 6 were combined during sequencing (due to low individual DNA content) and designated as passage 5.5. Thus, our training data included samples at passage 1 (N=2), passage 2 (N=1), passage 3 (N=2), passage 4 (N=2) and passage 5.5 (N=2).

Prior to training DNAmCULTURE, we sub-selected common CpGs between our MEF data, Petkovich et. al 2017 [42], and Thompson et. al 2018 [43] to generate a list of 28,323 common CpG sites (Supplemental Figure 2A). This was done so that our measure could be calculated in these external datasets to undergo a robust *in vivo* validation. Next, we conducted principal component analysis (PCA) using the training data and ∼28k sub-selected CpGs. PC1-4 captured greater than 70% of variance, with PC1 and PC2 exhibiting linear association with passage number (Supplemental Figures 1E-F). Based on our previous observations showing that combining PCA with elastic net yields more robust and reliable epigenetic age measures, we applied a similar strategy here [44,45]. Using all the components output from our PCA, we then combined this with a supervised machine learning approach. For instance, elastic net penalized regression was used to generate a predictor of passage number, but rather than feeding in CpGs as has been traditionally done in epigenetic clock development, we used PCs as predictors in our model. We have previously shown that this method is able to dramatically improve test re-test reliability and minimizes technical noise, while still capturing the critical signal from the data [44,45]. The lambda penalty was chosen via 10-fold cross-validation and resulted in a model that included six PCs (PC2, PC4, PC6, PC8, PC9 and PC29) (Supplemental Figure 2B-E). Overall, this measure is based on data from all 28,323 CpG sites, but is able to identify and combine the important patterns in genome-wide DNAm to generate a single score, DNAmCULTURE.

**Figure 2:**
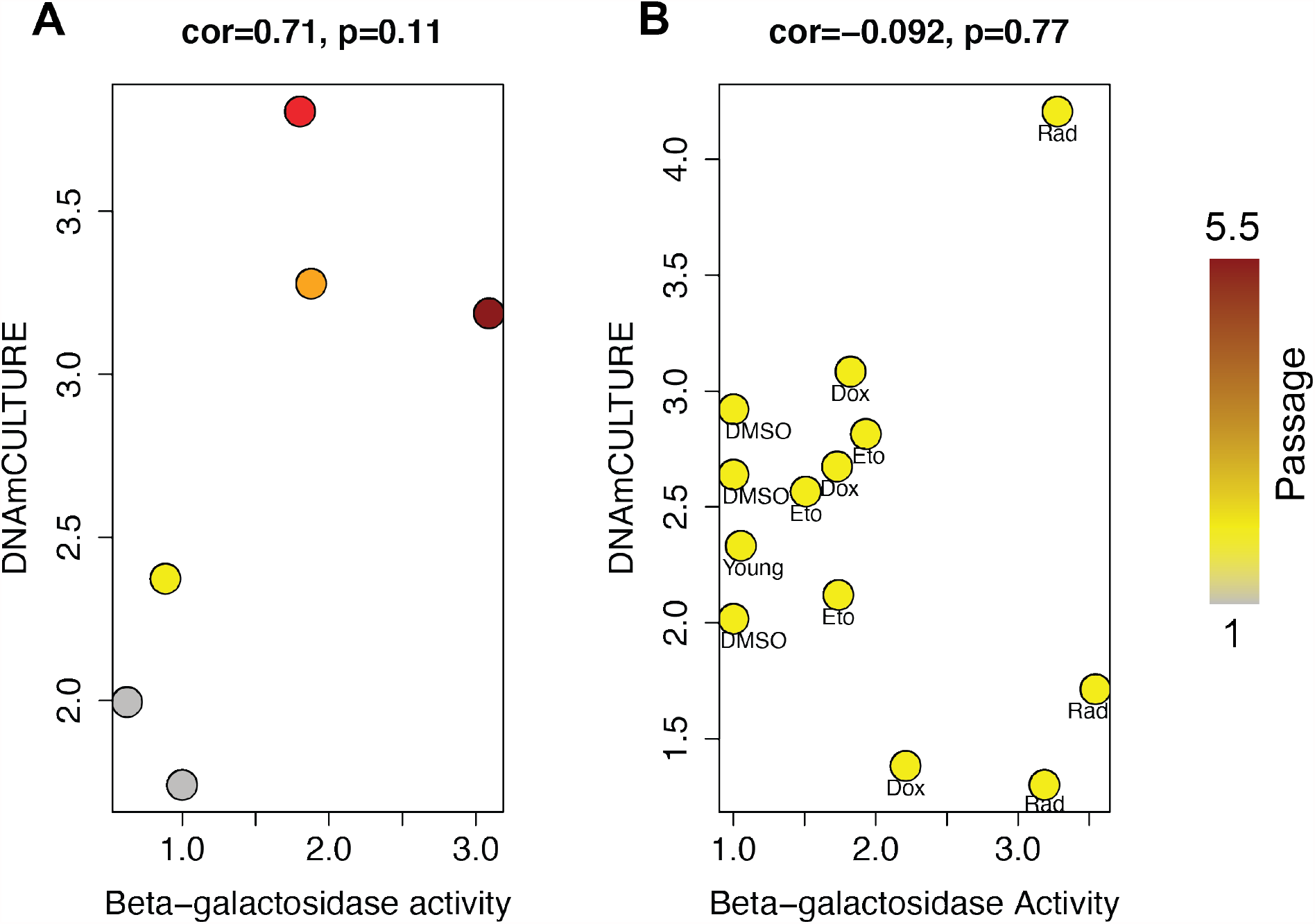
DNAmCULTURE fails to capture culture phenotypes independent of cellular passage. (A) DNAmCULTURE score in MEF3 replicative senescence validation samples, organized by Beta-Galactosidase (β-gal) activity. β-gal activity was determined by LogFITC fluorescence from C12FDG flow cytometry and normalizing geometric mean with negative samples (unstained) and dividing crude fluorescence by DMSO or young control, depending on experiment. β-gal activity calculation is further outlined in Supplemental Figure 1D. Note, grey=Passage 1, yellow=Passage 2, orange=Passage 3, red=Passage 4 and dark red=Passage 5.5. (B) DNAmCULTURE measured in irradiation and drug induced senescent samples against DMSO and young (Passage 2) controls. Note, Dox=Doxorubicin (1 µM), Eto=Etoposide (12.5 µM) and Rad=Irradiation (10 gy). Treatment occurred for 5 days. Senescence induction procedures are outlined in the methods. Passage independent experiments were conducted in MEF4-6 cell lines, which were validated by comparing passage 2 DNAmCULTURE score to MEF1 and MEF3 samples (Supplemental Figure 3C). β-gal correlations and statistical significance was determined using Pearson correlations.

Our results showed that DNAmCULTURE was highly correlated with passage number in both the training data (r=0.97), and in our independent validation samples (r=0.83), suggesting the marker is in fact progressively tracking with passage or time in culture (Figure 1C). In our training samples, we find that the measure shows a general linear increase. However, in the validation, there is a slight attenuation of the effect at the last passage. Given that we only have data on one sample at that passage, we cannot determine whether the non-linearity is real. One potential biological explanation is that there may be a deceleration at later cellular stages due to slowing in the growth rate from oxidative damage as cells approach or enter senescence.

### Distinguishing senescence from epigenetic aging

Replicative exhaustion in murine cells under normoxic (20% O2) conditions is a robust inducer of cellular senescence and we confirm in our study that MEFs arrest after 6 passages (Supplemental Figure 1B-D). Based on this, we tested whether our epigenetic measure was i) linked to senescence induction, likely as a result of chronic activity of a tumor repressor response to genotoxic stress, or ii) reflects aging changes that are independent of senescence state. When examining senescence-associated beta-galactosidase (SA-β-gal) as a function of DNAmCULTURE in passaged cells (Figure 2A), we observed a statistically insignificant association (r=0.71, p=0.11). Passaging (time in culture) and SA-β-gal activity are strongly associated, thus it is likely that the slight association between SA-β-gal activity and DNAmCULTURE is confounded by passage number. To test this further, we induced senescence in a passage-independent fashion using damaging dosages of irradiation (10 gy), doxorubicin (1 µM) and etoposide (12.5 µM). We show that each of these inducers elicits increased activity of SA-β-gal (Supplemental Figure 3A-B), however, SA-β-gal levels are not related to DNAmCULTURE (r=-0.092, p=0.77) (Figure 2B). Altogether, our results demonstrate that the extrinsic culture environment induces changes to the methylome in murine cells that is dependent on time in culture, rather than acute senescence.

**Figure 3:**
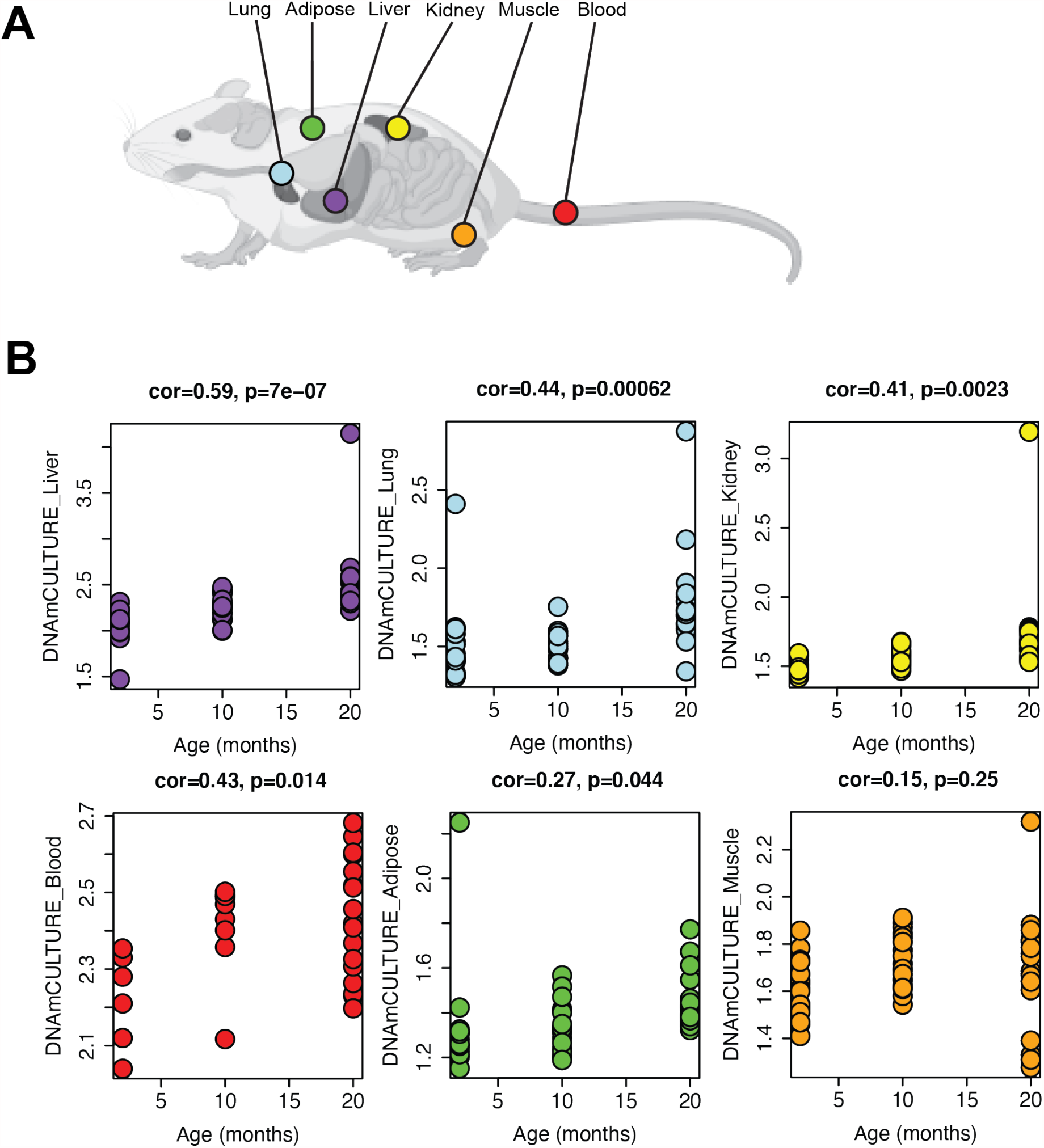
DNAmCULTURE recapitulates physiological aging in multiple tissues. (A) Longitudinal RRBS tissue data assayed from Thompson et. al 2018. (B) DNAmCULTURE measure determined in liver, lung, kidney, blood, adipose and muscle tissue at 2, 10 and 20 months in aged C57BL/6J mice. Age correlations and statistical significance was determined using Pearson correlations.

### *In vivo* validation and investigation into anti-aging therapies

Despite our ability to generate a measure that tracks *in vitro* aging, it is critical to test whether these changes mirror what is observed in aging tissues and cells *in vivo*. Thus, the robustness of DNAmCULTURE and its potential utility as a biomarker of *in vitro* aging was assessed using *in vivo* multi-tissue mouse DNAm data. When our measure was applied to data from Thompson et. al 2018—which included DNAm measured in multiple tissues at three timepoints (ages 2, 10 and 20 months) from C57BL/6J mice—we showed that DNAmCULTURE significantly increases with age in five of the six tissues: liver (r=0.59, p=7.0e-7), lung (r=0.44, p=6.2e-4), kidney (r=0.41, p=2.3e-3), blood (r=0.43, p=1.4e-2), and adipose tissue (r=0.27, p=4.4e-2) (Figure 3A-B). A moderate to low age increase was observed in skeletal muscle, although it was not significant (r=0.15, p=0.25). We next calculated DNAmCULTURE in a larger blood dataset from Petkovich et. al 2017 that included C57BL/6J mice between 20-1050 days of age and again observed a strong positive correlation with age (r=0.69, p=1.6e-23), adding further evidence that culture-derived alteration to the methylome are both physiologically relevant and widely represented in multiple tissue and cell types across the entire organism (Figure 4A).

**Figure 4:**
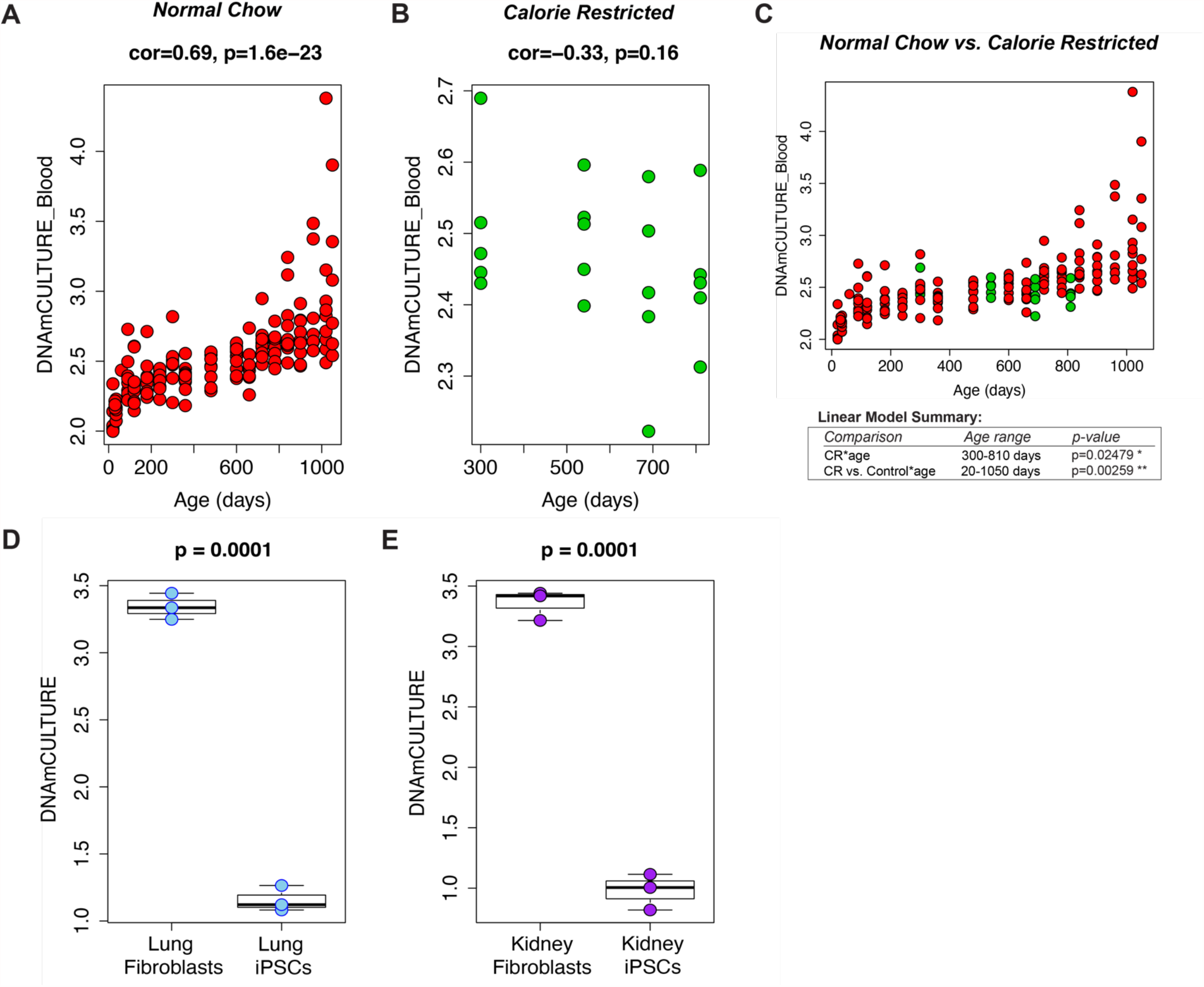
DNAmCULTURE predicts naïve culture states in caloric restricted mice and re-programmed fibroblasts. (A) DNAmCULTURE age-association determined from blood data assayed from a non-longitudinal cohort of aged C57BL/6J mice (20-1050 days) in Petkovich et. al 2017. (B) DNAmCULTURE age-association determined from blood data assayed from a calorie restricted (CR) cohort of aged C57BL/6J mice (300-810 days) in Petkovich et. al 2017. Calorie restricted mice began treatment at 14 weeks of age. Age correlations and statistical significance in (A) and (B) was determined using Pearson correlations. (C) Scatterplot demonstrating deceleration of culture aging in calorie restricted C57BL/6J mice, when comparing cohorts from (A) and (B). Red samples represent normal chow diet and green samples calorically restricted diet. Linear modeling demonstrates statistically significant deceleration in culture aging in CR samples (p=0.02479) as well as significant modulation in CR treated mice compared to normal chow controls (p=0.00259), when corrected by age. iPSC reprogramming in (D) Lung and (E) Kidney fibroblasts from Petkovich et. al 2017 demonstrates erasing of culture signature. Re-programming statistical significance calculations were determined via un-paired two-tailed t-test.

Using the Petkovich et. al data, we also found that DNAmCULTURE was responsive to dietary intervention (Figure 4B), such that calorically restricted (CR) mice exhibited significantly lower DNAmCULTURE scores relative to controls (p=0.00259), perhaps highlighting improved cellular maintenance and health from dietary intervention (Figure 4C). Finally, using the same dataset we showed that DNAmCULTURE also exhibits a decrease or re-setting in lung (Figure 4D) and kidney fibroblasts (Figure 4E) upon reprogramming to induced pluripotent stem cells (iPSCs) (p=0.0001).

### Clustering analysis confirms culture aging exists in physiological context and highlights Polycomb group (PcG) factors as important culture aging regulators

Given that DNAmCULTURE is a composite measure stemming from multiple aspects or domains of DNAm changes, we hypothesized that some of the signal it encompasses may be physiologically relevant, while others may be culture artifacts. For instance, we reasoned that supervised machine learning approaches, like elastic net, will prioritize strong signals in our culture models, despite whether they are physiologically relevant, limiting our ability to isolate important biological mechanisms. To address this, we applied consensus weighted gene correlation network analysis (WGCNA) to identify clusters (or modules) of highly co-methylated sites that are conserved across both *in vivo* [42,43] and *in vitro* data (Figure 5A, Supplemental Figure 4A). We identified 12 CpG modules, ranging in size from 105-678 CpGs. Most modules showed bimodal distribution in relation to distance from a transcription start sites (TSS), with many showing peaks at +/-100-1000 bp (Figure 5B).

**Figure 5:**
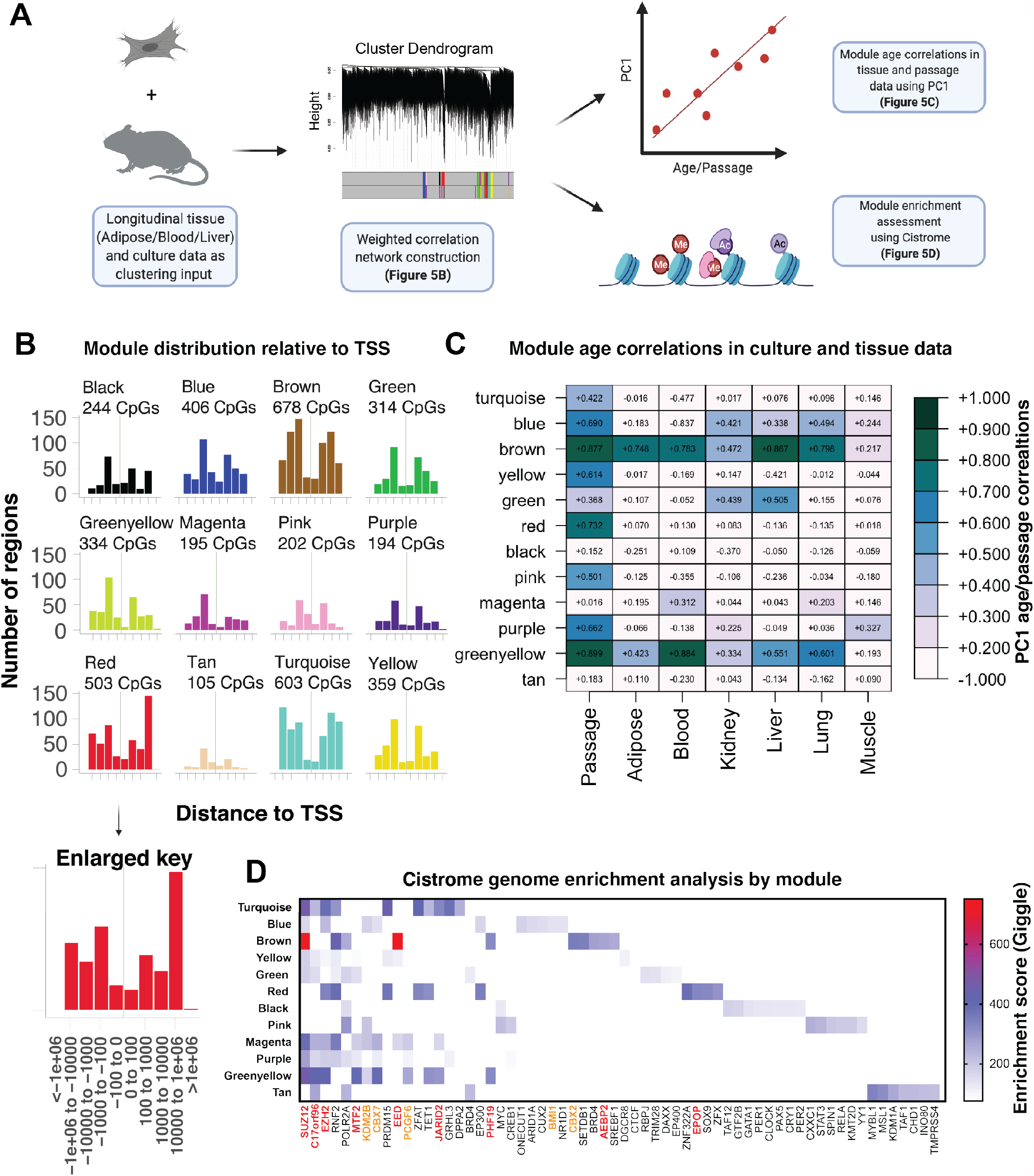
Clustering analysis confirms culture aging exists in physiological context and highlights Polycomb group (PcG) factors as important culture aging regulators. (A) Schematic outlining method of using longitudinal aging data (tissue + culturing) from Thompson et. al 2018 and the MEF1/MEF2 training data to cluster CpGs with WGCNA into distinct modules or ageotypes, which were then compared to *in vitro* passaging data and all tissues via principal component analysis and used to determine enriched genes using the Cistrome database. (B) Module distribution as determined by distance (per base pair) to transcription start site (TSS), generated using LolaWeb. Raw module CpGs were used to determine principal component correlations in (C), where kME selected CpGs were used to normalize enriched domains in (D), as further explained in Supplemental Figure 4B. (C) PC1 correlations of longitudinal tissue and MEF passaging data by module. (D) Module genome enrichment analysis using Cistrome database from 100 CpG input selected by kME. Enriched genes were further normalized by randomly selecting 100 CpGs from the background 27,035 CpGs used to create the modules and correcting each enriched GSM_IDs Giggle score. Note, the enrichment analysis is displaying the top normalized GSM_ID Giggle score for each enriched gene module relationship. Enriched genes are sorted by decreasing module frequency. Giggle score represents a rank of significance between genomic loci shared between query file and thousands of genome files from databases like ENCODE. Red genes = PRC2 complex or mediator, Orange genes = PRC1 complex or mediator and Black genes = non-Polycomb related genes.

Next, we estimate module eigengenes and tested their associations with passage number (*in vitro* MEF data) and age (*in vivo* tissue data). Eigengenes were calculated as PC1 estimated from the *in vitro* data and then applied as validation to the *in vivo* data (Figure 5C). Using these values, we observed several modules that appear to be artifacts of *in vitro* aging (turquoise/yellow/red/pink/purple), such that they showed progression with passage number in MEFs, but did not track with age in tissues. However, two modules (brown and greenyellow) stood out as being potentially translational (culture and tissue). For instance, the brown module was strongly correlated with passage number (r=0.88), as well as age in liver (r=0.87), lung (r=0.80), blood (r=0.78), adipose (r=0.75). It was also moderately correlated with age in kidney (r=0.47) and weakly correlated with age in skeletal muscle (r=0.22). The greenyellow module exhibited strong correlations with both passage number *in vitro* (r=0.90) and age in blood (r=0.88), while showing moderate age correlations with lung (r=0.60), liver (r=0.55), adipose (r=0.42), and kidney (r=0.33), and a weak correlation with age in skeletal muscle (r=0.19).

Finally, to garner more biological insight into potential mechanisms at work in conserved modules, we assessed genome enrichment of transcription factor (TF) binding motifs and chromatin regulators using the Cistrome database. This was done by comparing each module by TF and chromatin regulator enrichment score (giggle score). The giggle score represents a rank of significance between genomic loci shared between query file and thousands of genome files from databases like ENCODE. Given that scores tend to increase for lists with a greater number of input genomic locations (and thus would be biased by module size), we normalized each module prior to determining the enrichment score so that only 100 CpG locations were being assessed for each module. For instance, we selected the top 100 CpGs with the highest kME values in a given module. kME is estimated as the correlation between CpG values and the module eigengene and can be used to infer connectivity or identify “hubs” of a module. For the background CpGs, we selected 100 CpGs from the 27,035 CpG background at random and used the background giggle score to blank any hit overlap from the modules. The final 100 input CpGs for each module are reported by genomic partition distribution (Supplemental Figure 4B) and scatterplots of each raw Cistrome distribution are reported by module (Supplemental Figure 4C). We compared the 10 top GSM_IDs (query datasets) from each module to determine the most enriched gene regulators.

The Cistrome analysis (Figure 5D) reveals the novel finding that Polycomb repressive complex 1 and 2 (PRC1 and PRC2) networks are highly enriched in translational modules (brown and greenyellow), highlighting PcGs as key epigenetic regulators in both culture and physiological aging. With almost all of the top hits for greenyellow (9/10) occurring in PcGs and the highest giggle enrichment scores occurring in SUZ12 and EED (PRC2 components) for the brown module, our data suggests PcGs regulate physiologically relevant culture aging phenotypes.

## Discussion

Consistent evidence supports the notion that DNA methylation is dramatically altered with aging [26]. While numerous epigenetic clocks have been derived approximating these changes using tissue samples from a variety of mammalian species, the exact drivers of epigenetic aging are unknown [26-30]. Likewise, it is unclear how the methylome progresses in an artificial aging context, such as in cellular culture. Given that well characterized culture systems exist [35], we aimed to classify potential epigenetic drivers of culture aging and determine if such changes recapitulate physiological aging in various tissues and biofluids. We rationalized that with the widespread use of culture models throughout biology and medicine, many fields would greatly benefit from clarifying the underlying epigenetic phenotypes that exist in culture and whether relevant markers of cellular dysfunction can be trained for use in accelerating mechanistic and drug development discoveries.

By exhaustively passaging primary MEFs under normoxic conditions (20% O2), we trained a DNAm predictor of passage number (time in culture), called DNAmCULTURE and demonstrate that it not only accurately tracks passage number (Figure 1C), but also strongly correlates with age in multiple tissues (Figure 3A-B, Figure 4A), is modifiable by dietary intervention (Figure 4B-C), and exhibits resetting upon reprogramming to pluripotency (Figure 4D-E). For instance, we found that DNAmCULTURE measured in whole blood of C57BL/6 mice was correlated with age (Figure 4A), despite being trained using cultured fibroblasts. Similarly, DNAmCULTURE exhibited age-related increases in liver, lung, kidney, blood and adipose tissue. Interestingly, it did not strongly correlate with age in skeletal muscle (Figure 3B), which may reflect the fact that skeletal muscle remains mostly postmitotic in adulthood. The link between proliferation and DNAmCULTURE was also observed when comparing the other tissue types. For example, we observed differences in both age correlation/slope, and in the absolute scores when comparing tissues. Overall, samples from liver and blood appeared to exhibit the highest values (Figure 3B), which may reflect the higher proliferative capacity of cells in these samples or the renewable nature of both hepatocytes and blood cells, perhaps suggesting that lifetime damage is somehow cataloged by the methylome [46-48]. This is also substantiated by the observations that epigenetic aging is not linear with time [26]. For instance, previous epigenetic clocks have been shown to increase rapidly during development and then decelerate after full maturity. We were able to observe this same trend in our data. We found that DNAmCULTURE exhibited a sigmoidal function with age, characterized by accelerated aging during development, a slower and more linear increase after about 150 days, and exponential increases at late life (Figure 4A).

Despite the evidence of a relationship between replication and epigenetic aging, our data suggests that this is independent of senescence accumulation. For instance, we showed that drug and irradiation induced senescence in MEFs was not associated with changes in DNAmCULTURE. Further, replicative senescent MEFs were non-significantly associated (Figure 2A-B), suggesting senescence states are related to DNAmCULTURE, but only in a passage dependent fashion. One possible explanation is that DNAmCULTURE is capturing chronic epigenetic stress from genotoxic damage experienced throughout extended time in culture, rather than acute damage. Previous studies provide evidence that MEFs are highly sensitive to genotoxic stress, especially when cultured under normoxic conditions (20% O2) [35,39]. The consensus in the field is that when MEFs experience much higher O2 tension than what they experience *in vivo* (3% O2), they undergo accelerated oxidative damage and after 5-7 passages they initiate replicative senescence programs [35,39]. DNAmCULTURE sheds light on the possibility of cell extrinsic factors as drivers of epigenetic aging. A number of previous studies provide evidence of this by suggesting that intrinsic drivers, like telomere attrition, are unlikely to drive culture phenotypes in accelerated aging murine models, as mice have exceptionally long telomeres and only become growth arrested in a telomere dependent fashion from loss or deletion of the telomerase RNA component (mTR) and p53 [31,49-53].

The potential links between epigenetic aging, replication, and genotoxic stress may also explain the age-related increase in cancer susceptibility, particularly among highly proliferative tissues and cells. For instance, we and others have previously reported that epigenetic age changes are also observed at increasing rates in tumors and/or the normal (or non-afflicted) tissues of individuals with cancer. We reason that the epigenetic changes captured by measures like DNAmCULTURE may underlie susceptibility to spontaneous transformation or oncogenicity [54]. Cells that eventually evade senescence from mutational events may promote oncogenic states, allowing continued mitotic events and increased damage accumulation, as a function of cell turnover [35]. In moving forward, it will be critical to utilize future *in vitro* experiments to determine the mechanisms driving epigenetic changes as a function of either mitotic rate (replication “ticking”) and/or prolonged exposure to genotoxic stress.

While substantial work has gone into developing biomarkers than enable researchers to track aging changes *in vivo* and *in vitro*, the ultimate goal is to develop measures that are also modifiable to intervention. Using DNAm assessed in blood, we reported the effects of two promising interventions in aging— caloric restriction (CR) and cellular reprogramming. Our results suggested that DNAmCULTURE showed strong response to CR when assessed in blood (Figure 4C). Multiple studies suggest that CR acts by reducing DNA damage accumulation and mutations that progress with age [55], where others show CR downregulates key growth hubs like the insulin/IGF1 pathway [56]. Importantly, IGF1 is a growth factor that stimulates cell proliferation and can promote cancer via inhibition of apoptosis [57,58]. Interestingly, CR, without malnutrition, has also been shown to reduce cancer incidence and progression in mice [59,60]. Our results suggest that CR could be acting via the epigenome to regulate DNA damage maintenance by slowing cellular turnover and thus damaged states, or perhaps from enhanced DNA repair. Additionally, our results showed that the longer mice underwent CR, the more they diverged from normal controls on the basis of DNAmCULTURE. This could suggest that prolonged CR does not simply reverse or retard epigenetic aging momentarily, but actually decelerates the rate of change with age.

We also report renewal in lung and kidney fibroblasts indicative of naïve culture states following reprogramming to iPSCs, supporting the conclusion that DNAmCULTURE can not only be slowed, but actually reversed (Figure 4D-E). For instance, both lung and kidney fibroblasts were predicted to be equivalent to cells passaged just over 3 times, while all iPSC derivatives were predicted to approximate cells at passage 1. This suggests that the major epigenetic changes acquired during culturing and/or tissue aging can be reset to some extent. It is unlikely DNA damage and the resulting genome instability is reversible, thus we propose that DNAmCULTURE may be capturing transient epigenetic programs that control survival, proliferation, and cellular maintenance. In moving forward, it will be critical to establish how the epigenome is remodeled during full or partial reprogramming. It may also be the case that this rejuvenation phenomena is driven by distinct types of epigenetic changes that share specific functional characteristics. Although we and others have demonstrated that classical epigenetic clocks (trained on tissues) can capture culture aging changes like reprogramming, it is difficult to parse out the functionally relevant mechanisms from the “black box” algorithms [44,61]. Moreover, we have previously shown that epigenetic clocks represent composites of distinct types of epigenetic phenomena that may not share the same mechanistic underpinnings. As such, it is difficult to experimentally test the causes and consequences of these clocks, thus limiting their utility as experimental tools in translational research.

In the current study, we also tested whether we could distinguish different “types” of DNAm changes in our data. To identify the methylation changes in culture aging systems that are the most physiologically relevant to aging in tissues, we applied a network-based approaches to group CpGs into modules and then assess their relative patterns across datasets. Our results clearly demonstrate that *in vitro* DNAm changes captured by the red, yellow and pink modules were not physiologically relevant, suggesting that they may be reflective of culturing or MEF-specific artifacts. In contrast, CpGs in the brown and greenyellow modules appear to capture a common epigenetic aging phenotype that is established in both physiological and culture aging context (Figure 5A-C). To better understand the molecular underpinnings of CpGs in these two modules, we used genome enrichment analysis to determine potential drivers or regulatory features.

By utilizing the Cistrome database we provide evidence that PcG factors, including both PRC1 and PRC2, are key factors in physiologically-relevant culture aging (Figure 5D). It is well established that the tri-methylated histone H3 at lysine 27 (H3K27me3) mark denotes transcriptional silencing with PRC2 involved in early development and PRC1 later during aging as the more active maintenance factor [62]. The catalytic subunit of PRC2, EZH2, is routinely overexpressed in oncogenesis [63], promoting uncontrolled cell growth, as many repressed downstream genes of H3K27me3 are tumor suppressors [64], but the role of PRC2 and its domains are conflicted in aging. In certain species and cell types, EZH2 mutations reduce H3K27me3 and confer longevity [65,66], although in others reduction of H3K27me3 is associated with aging [67]. The relationship between the catalytic subunit (EZH2) and its co-factors SUZ12, EED, RbAp48 and AEBP2, which are highly involved with allosteric recognition and binding of substrates like *S*-Adenosyl methionine (SAM), is multi-factorial, with many opportunities for perturbations. As an example, multiple studies demonstrate EZH2, SUZ12 and EED are essential components for proper functioning, but RbAp48 and AEBP2 are not [68,69]. Our reported translational modules (brown/greenyellow) further support the notion that PcGs are important aging factors.

## Conclusion

We report a novel mouse epigenetic measure of culture aging, termed DNAmCULTURE, that is able to recapitulate epigenetic changes observed in multiple *in vivo* tissues. We confirm that DNAmCULTURE is independent of senescent state, and instead appears to capture progressive cellular changes that may confer susceptibility to senescence and/or tumorigenesis. We also provide evidence for potential modifiability in the form of deceleration as a function of CR or reprogramming. Finally, our results implicate DNAm changes that may be functionally related to Polycomb group (PcG) factors like EED and SUZ12. Overall, this study demonstrates that physiologically relevant DNAm changes can be modeled *in vitro*, which in the future can be used to interrogate mechanisms involved in epigenetic aging and/or facilitate *in vivo* aging discoveries.

## Methods

### Experimental

#### Mouse embryonic fibroblast extraction

Mouse embryonic fibroblasts (MEFs) were harvested at day 12.5 of gestation. Two females were used. From the first female, 9 embryos were sacrificed and split into three cell lines, MEF1-3 From the second female, 10 embryos were sacrificed and split into three cell lines, MEF4-6.

Extraction was achieved by separating embryos into separate wells in a 6 well dish using PBS, removing inner embryo and using forceps to carefully remove limbs, head and internal organs from dorsal region. The dorsal region was then cut and trypsinized for 10 minutes at 37°C. To quench reaction cells were transferred to a 15 mL falcon tube and spun for 3 minutes at 1200 rpm, then supernatant was aspirated and resuspended with 10 mL DMEM. P0 cells were split once to expand cell number prior to freezing. Approximately 2 mL of cells were incubated overnight with 8 mL DMEM and following growth were trypsinized and either passaged for experiments or stored at −80°C in DMEM/DMSO (90:10).

#### Replicative passaging and cell culture

Cells were split/passaged 6 times, where flow cytometry/confocal microscopy and RRBS sequencing were conducted at each passage.

Cells were split according to the following seeding density -p100 −0.5×10^6^. cells, p60 −0.25×10^6^. cells and 6 well −0.125×10^6^. cells -and were counted using an Invitrogen countess and cell counting chamber slide with trypan blue. For media, we used DMEM + 10% FBS + 1% PENSTREP. Note, later passaged cells had a lower platting efficiency when inspected visually 24 hours after seeding, thus we used a cell scraper prior to transfer otherwise senescent cells remained attached to the dish. Cells were split at approximately 95% confluence which occurred around 3-4 days in P1-3 and 5-8 days in P4-6.

#### Beta-galactosidase flow cytometry and confocal microscopy

To conduct beta-galactosidase flow cytometry, approximately 0.25×10^6^. cells were seeded into p60 dishes and pre-treatment was conducted approximately 16 hours after seeding. Cells were first pre-treated with Bafilomycin A1 (Selleckchem: S1413, 622.83 g/mol, 100 µM stock). Existing DMEM was aspirated, then cells were washed with PBS and replaced with treated Bafilomycin A1 DMEM for 30 minutes at a final concentration of 100 nM. Following Bafilomycin A1 pre-treatment to normalize lysosome activity, C12FDG (Invitrogen: D2893, 853.92 g/mol, 10 mM stock) was added directly to the existing media for 90 minutes at a final concentration of 20 µM. Note, due to light sensitivity, exchange was conducted in a dark environment.

For determining beta-galactosidase activity via flow cytometry, treated cells were trypsinized (1 mL-p60) for 5 minutes at 37°C and then quenched using 3 mL DMEM. Note, cells were completely detached using a cell scraper prior to transfer otherwise senescent cells remained attached to the dish. After thorough resuspension, cells were transferred directly to a filter top tube and spun for 3 minutes at 1200 rpm. Supernatant was aspirated and cells were resuspended in 100 µL PBS and immediately assayed using a 488 nM laser on a StratedigmS1000 benchtop flow cytometer. Fluorescence intensity was normalized and baselined using an unstained sample. FlowJo (10.6.1) was used to analyze data. Beta-galactosidase activity/senescence activity was determined as LogFITC treated geometric mean/control geometric mean after normalizing to untreated control.

For determining beta-galactosidase activity via confocal microscopy, cells were split into 12 well dishes with a glass cover slide at the bottom of each well. Following Bafilomycin A1 and C12FDG treatment, media was aspirated and cells were washed with PSB 3x, fixed with 4% PFA/PBS (10 minutes), followed by 2x PSB washes and then counter stained with DAPI and mounted onto cover slips. Fixed cells were immediately imaged at 4x, 10x and 40x resolution using a Keyence confocal cytometer.

#### Senescence induction

We induced senescence using previously established conditions [48]. In brief, MEFs were thawed and allow to expand for one passage, then split to a normalized seeding density of 0.25×10^6^. cell/p60 and 0.125×10^6^. cells/6-well and treatment was conducted for 5 days. Note, senescence induction experiments were conducted at passage 2. Doxyrubicin (Sigma: D1515, 1 µM), Paclitaxel (Sigma: T7402, 50 nM) and Etoposide (Sigma: E1383, 12.5 µM) were all dosed into DMEM when the cells were split and media was not replaced for the duration of the 5-day treatment. We irradiated cells (10 Gy) using cesium irradiation and collected these cells after 5 days as well.

#### DNA preparation and quantification

DNA was extracted from selected samples prior to RRBS sequencing using a Qiagen DNeasy Blood and Tissue extraction kit (69504). Note, samples were treated with proteinase K and RNAse A and eluted with 200 µl elution buffer. Following final elution, DNA was verified using nanodrop (Thermo Scientific). Spin concentration was used as necessary with low DNA content samples. Prior to library preparation we used a qubit fluorometer (Thermo Scientific) to quantify the extracted genomic DNA. All samples were assigned a single-blinded code and randomized for library preparation and sequencing to control for any batch errors.

#### Library preparation and reduced representation bisulfide sequencing

Library preparation was conducted using EZ DNA Methylation RRBS Library Prep Kit (Zymo: D5461), according to manufacturer’s recommendations. Randomized and pooled samples were sequenced on four Illumina NovaSeq6000 SP lanes (100 bases single-end mode). Note, each lane produced more than 400M reads.

### Statistical Analysis

#### Data preprocessing

FastQC (v0.11.8) was used to assess the quality of the raw reads and adapter-trimmed reads (cutadapt, version 2.5). Reads were mapped to the GRCm38 RRBS genome using BSBolt v0.1.2 (https://github.com/NuttyLogic/BSBolt). Methylation was called and the CpG methylation matrix was assembled for CpG sites common to all samples and covered by more than 10 reads. The final matrix consisted of 474,128 CpG sites.

#### *Training and Validation of* DNAmCULTURE

R was the primary platform used for statistical analysis. After selecting overlapped CpGs between training and all validation studies, PCA (without scaling) was conducted in the training sample. Training using PC components was conducted as described previously [44,45]. PCs were then fed-into a penalized elastic net regression as variables to train a predictor of passage number, called DNAmCULTURE. Lambda penalty represented the value with lowest mean-squared error, selected via 10-fold cross-validation.

To validate the measure, PCs were estimated in external datasets using the loading from the training sample. These PCs were then incorporated into the selected elastic net model. Pearson correlations were used to assess associations between DNAmCULTURE and 1) passage number in both the training and validation sample, 2) β-gal activity in senescence induced MEFs, and 3) age in multi-tissue in vivo samples. Two-tailed t-tests were used to compare significance in iPSC reprogramming and in MEF4 validation. To test for associations with CR, OLS regression was used that included age, CR and an interaction term (age*CR).

#### WGCNA and module construction

Consensus WGCNA [70] was conducted using four input datasets—MEF training samples (replicates 1 and 2), and the Thompson et al. data for blood, liver, and adipose. The remaining Thompson et al. data (kidney, lung, muscle) was deliberately excluded from WGCNA so as to have a true validation. Adjacency was estimated for each dataset based on biweight midcorrelations and negative correlations were treated as unconnected in the network (signed network). Adjacencies were then converted to Topological Overlap Matrices (TOMs) and combined into a single consensus TOM, such that overlap for each CpG pair was designated as the minimum dissimilarity score across the four individual TOMs. Hierarchical clustering was then conducted with the following parameters: deepSplit=1, cutHeight=0.95, minClusterSize=50, and distance=(1-consensus TOM), method=“average). This resulted in a network with n=16 modules. Given that similar modules can often be split by WGCNA, we next tested whether modules should be merged. This was done by estimating module eigengenes and then assessing dissimilarity between modules. Using a cut height of 0.4, the 16 modules were merged into 13 that served as our final modules for all remaining analyses. One feature of WGCNA is the ability to estimate module eigengenes, which serve as single quantitative value meant to represent the core signal of a whole module—that can consist of tens to thousands of individual variables. Typically, PC1 from PCA run on all variables in a module is used to represent the module eigengene. However, the traditional WGCNA package estimates this separately for all dataset meaning that the eigengenes may not be based on the same equations across datasets (variables can have different loadings). This may cause a bias in results and make validation less straight forward. To overcome this, we estimated PC1 for each module using the MEF training data and then applied these loading to all other datasets, including those used in WGCNA and thus that were held-out. Finally, we tested whether the module eigengene values were associated with either passage number (MEF data) or age (multi-tissue data).

#### Cistrome genome enrichment analysis

We used the Cistrome gene analysis tool kit (http://dbtoolkit.cistrome.org/) to determine enriched genes. We selected the top 1k hits and used the mm10 reference. The outcome of the enrichment analysis was reported as a Giggle score, which is a rank of genome significance between the input file and thousands of genome files from databases like ENCODE. It is important to note, that Cistrome is constantly updating genome files, thus the enrichment analysis was conducted at the same time. Additionally, we selected 100 CpGs from each module using kME to select the most central 100 CpGs. Sub-selected CpGs are reported via genomic partition in Supplemental Figure 7A. For selecting the background 100 CpGs we randomly selected the 100 CpGs from the cohort of 27,035 CpGs. For giggle score reporting, we plotted the raw giggle score of each resulting module query, although any file (GSM_ID) that was also a background hit was corrected using the formula; GSM_ID_Hit-GSM_ID_Background=GSM_ID_Actual. Note, when the background GSM_ID was not present there was no correction. We report raw giggle scores in a scatterplot format in Supplemental Figure 7B and the corrected values (Top 10) in Figure 5D.

Genomic partitioning and CpG locations were determined using LolaWeb (http://lolaweb.databio.org/)

## iv. Acknowledgments

This work was funded by support by the Glenn Foundation (award for Research in Biological Mechanisms of Aging) and the National Institute on Aging (R01AG068285 and R01AG065403).

## vi. Authors contributions

Contribution is based on authorship.

## vii Data availability

The data that support the findings of this study are available from the corresponding author upon reasonable request.

## ix. Tables

None to report.

## x. Figures

**Supplemental Figure 1:**
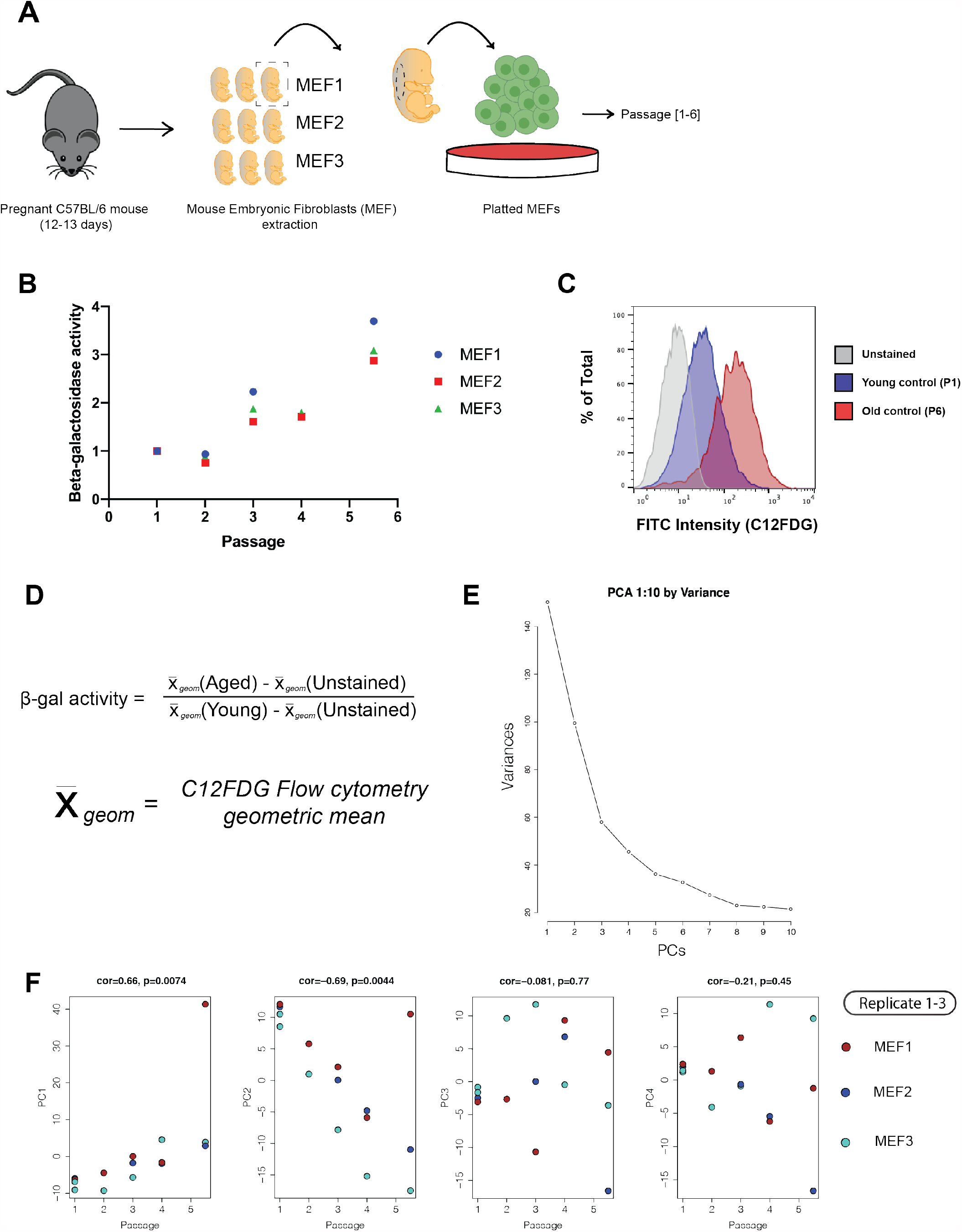
Mouse embryonic fibroblast extraction, passaging and validation. (A) Schematic demonstrating MEF extraction, illustrating embryos were dorsally derived from 12.5-day gestation C57BL/6 mice, then passaged 6x. Note that each biological replicate was composed of 3 embryos. (B) Plot of flow cytometry data demonstrating increased Beta-galactosidase (β-gal) activity with passage, measured by FITC fluorescence from C12FDG and normalizing geometric mean with negative samples (unstained) and dividing crude fluorescence by young control (Passage 1). (C) Representative flow cytometry plot demonstrating older MEFs have greater beta-galactosidase activity. (D) β-gal activity calculation. (E) Plot measuring PCA components [PC1-10] from MEF1-3 (passages 1-6) as a function of variance. (F) Principal component analysis of all MEF cell lines used for training (MEF1-2) and validating (MEF3) DNAmCULTURE.

**Supplemental Figure 2:**
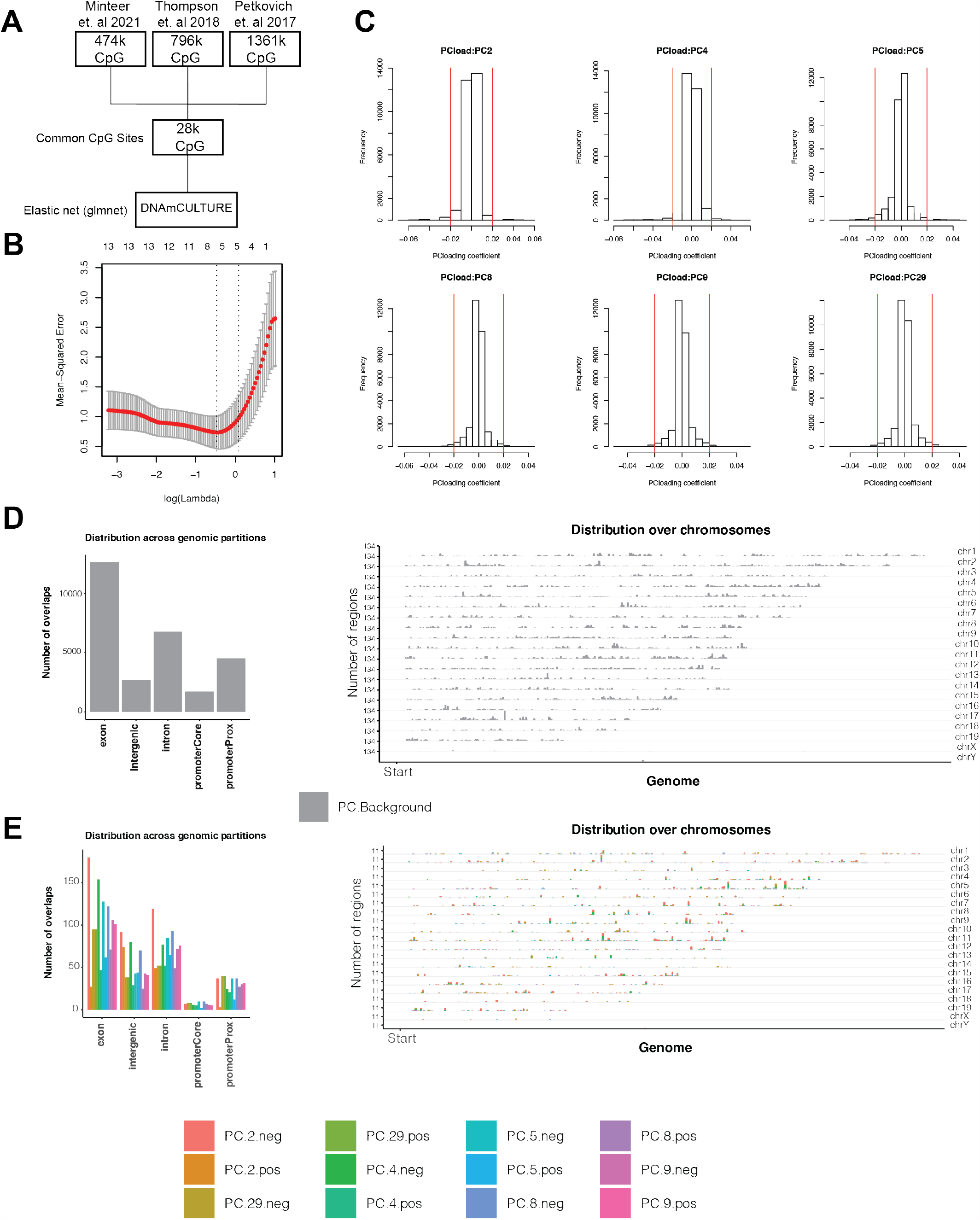
DNAmCULTURE construction, PC loading and CpG distribution. (A) Common CpGs (28,323) between MEF experimental data, Petkovich et. al 2017 and Thompson et. al 2018. (B) Elastic net penalized regression plot generating lambda minimum for selecting PCs (PC2, PC4, PC5, PC8, PC9 and PC29) for DNAmCULTURE. (C) Histogram of loaded PCs in DNAmCULTURE, plotted by PC loading coefficient and frequency of total 28,323 CpGs in measure. Red abline represents 0.02 cutoff used to determine CpG drivers of DNAmCULTURE. (D) CpG distribution across chromosomes and genomic partitioning of raw 28,323 CpGs used in DNAmCULTURE, generated by LolaWeb. (E) CpG distribution across chromosomes and genomic partitioning of CpG drivers (N=3087), as determined by 0.02 loading coefficient cutoff, also generated by LolaWeb.

**Supplemental Figure 3:**
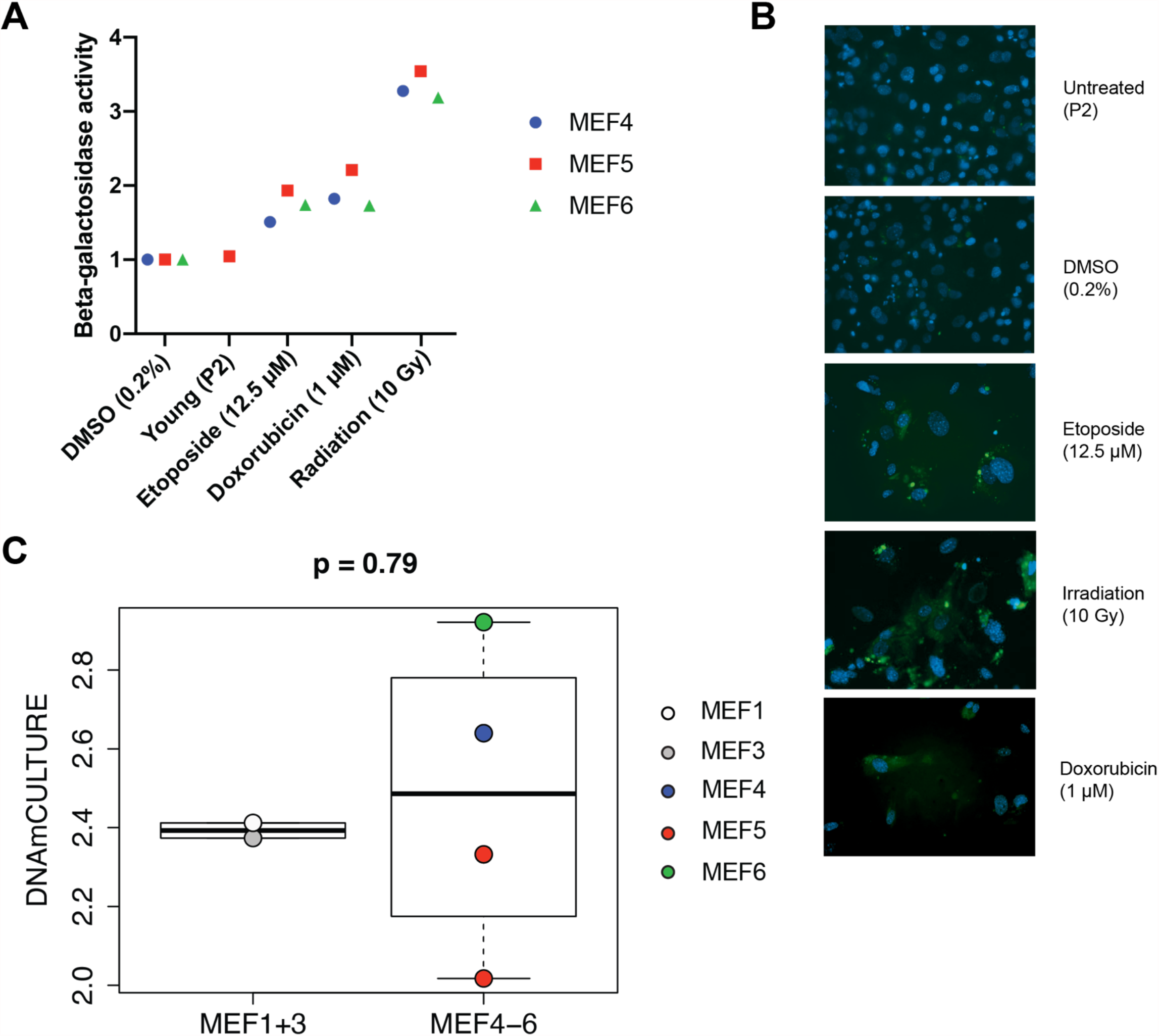
Passage independent senescence induction and MEF technical replicate validation. (A) Beta-galactosidase activity measured in MEF4-6 with various damaging and control conditions. β-gal activity was determined by LogFITC fluorescence from C12FDG flow cytometry and normalizing geometric mean with negative samples (unstained) and dividing crude fluorescence by DMSO or young control, depending on experiment. β-gal activity calculations are further explained in Supplemental Figure 1D. (B) Representative confocal microscopy images (40X) using C12FDG (green) and counterstained with DAPI (blue), confirming senescence is achieved from irradiation and drug treatment (doxorubicin and etoposide). (C) DNAmCULTURE measured in all MEF replicates for passage 2 samples under control conditions (either Young untreated or DMSO), demonstrating no significant inherent variation exist between replicates. Statistical significance calculations were determined via un-paired two-tailed t-test.

**Supplemental Figure 4:**
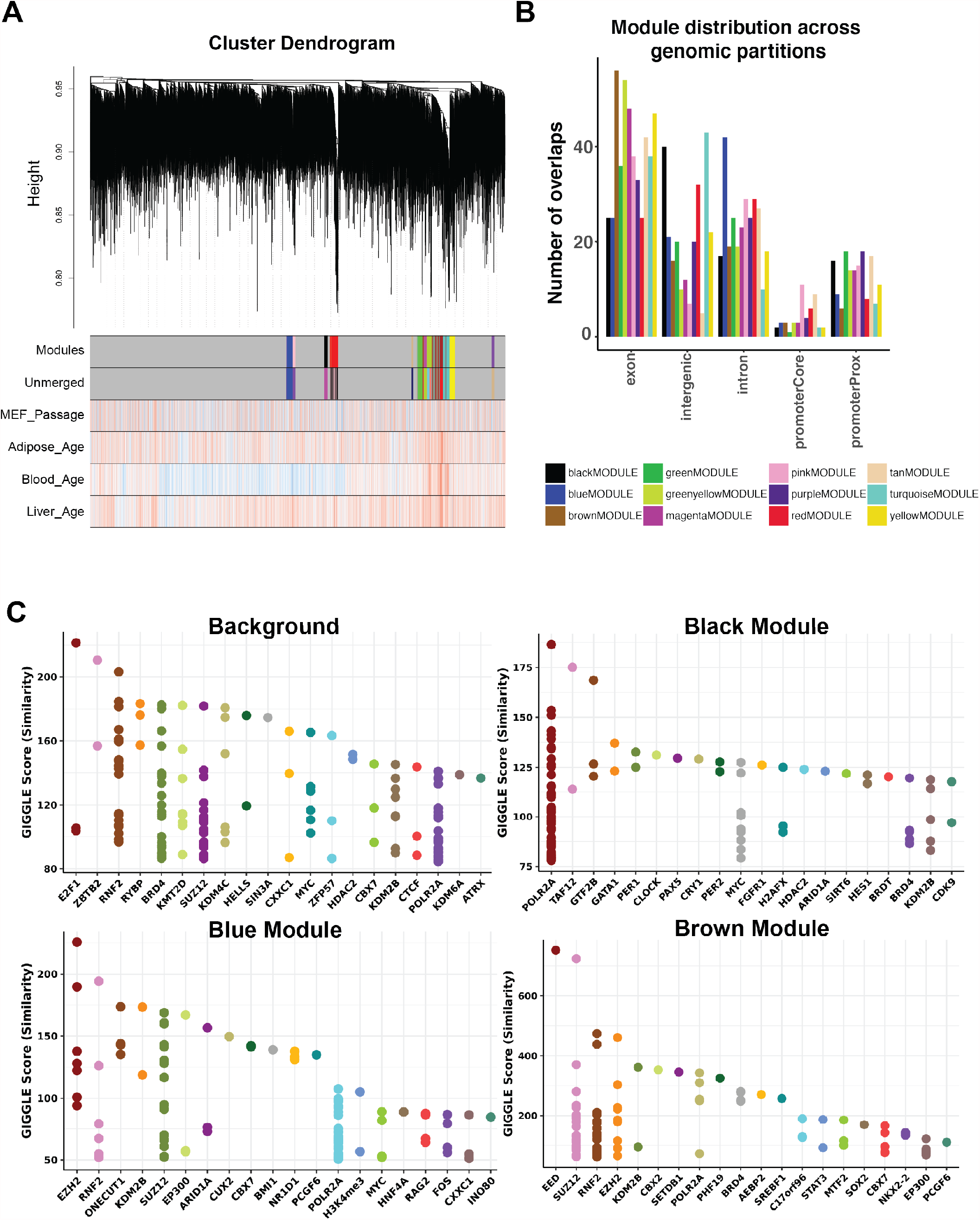
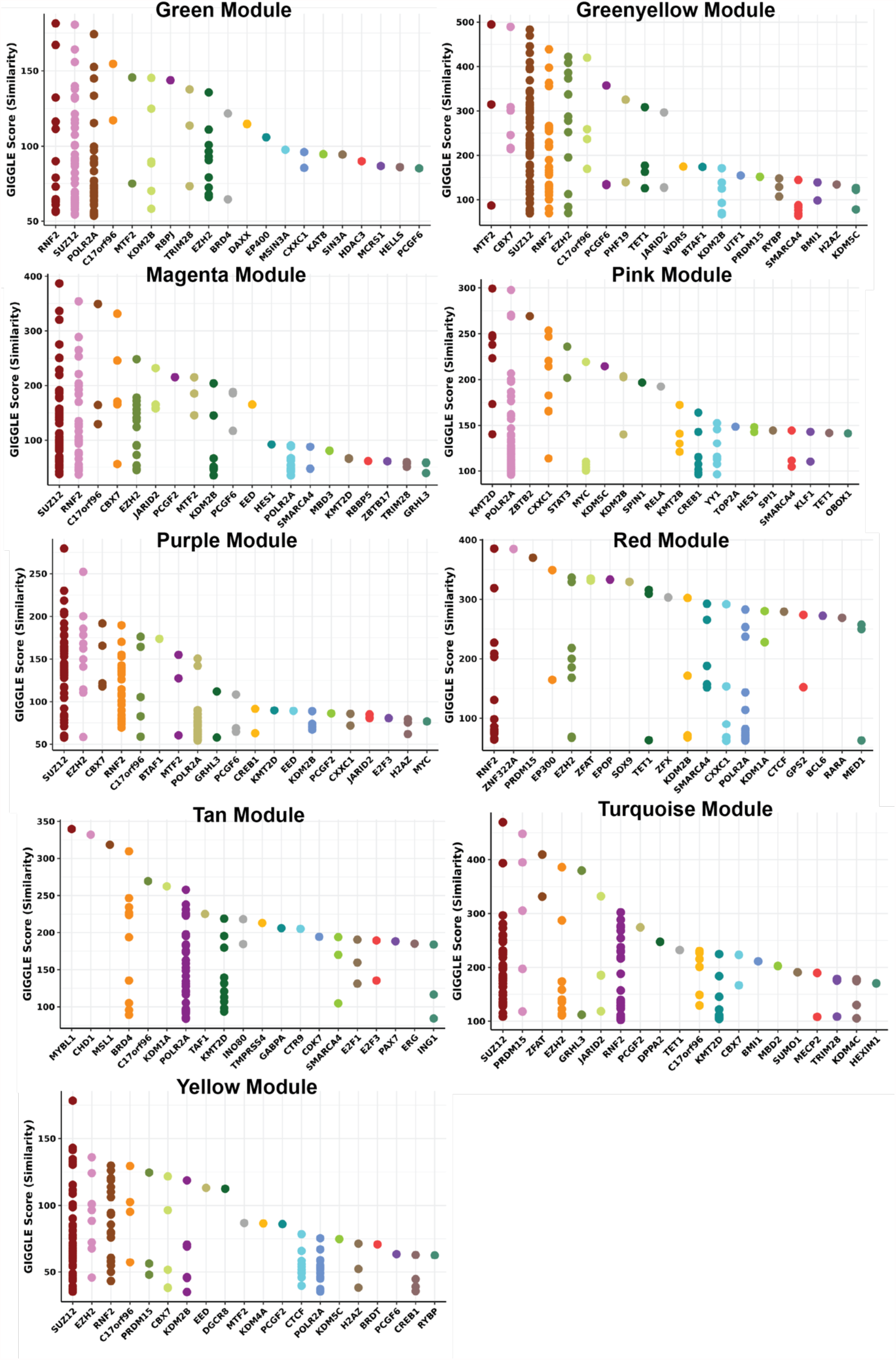
Network construction and Cistrome enrichment analysis. (A) Cluster dendrogram demonstrating 12 modules and age/passage associations with culture, adipose, blood and liver input data. (B) Genomic partition (generated by LolaWeb) of top 100 CpGs, as determined by the most central CpGs by kME, of the 12 selected WGCNA modules. kME selected CpGs were used to normalize enriched domains in (C). (C) Scatterplots of top 20 enriched genes in each module, as determined by Cistrome, prior to any background baselining. Note, the top 100 CpGs, as determined by kME, were used as the input for each query in order to cross compare modules by Giggle score. 100 CpGs were selected at random from the 27,035 background CpGs that were used as the input for clustering analysis. Giggle score represents a rank of significance between genomic loci shared between query file and thousands of genome files from databases like ENCODE.

## xi. Appendices

None to report.

